# Validation of tWO novel primers for THE promising amplification of the mitogenomic Cytochrome c Oxidase subunit I (*COI*) barcoding region in *Artemia* aff. *sinica* (Branchiopoda, Anostraca)

**DOI:** 10.1101/2022.05.07.491003

**Authors:** Alireza Asem, Chun-Zheng Fu, Ning Yang, Amin Eimanifar, Ying Cao, Pei-Zheng Wang, Chun-Yang Shen

## Abstract

Due to the lack of a taxonomic key for the identification of *Artemia* species, molecular markers have been increasingly used for phylogenetic studies. The mt*COI* marker is a regularly considered marker for the molecular systematics of *Artemia* populations. The proposed universal and specific primers have mostly failed to amplify the *Artemia* aff. *sinica* mt*COI* marker, and on the whole, the successfully amplified products of the PCR were inefficient, primarily through the representation of poly-peak or incomplete sequences. We presumed that if a forward primer could be developed regarding the joint regions of the last part of the previous gene (tRNA_*Tyr*_) and the beginning of the target gene mt*COI*, the sequence could be relevant to the target-sequence of mt*COI*. Thus, here, we describe a new set of primers, which could be used to amplify the mt*COI* barcoding region of *Artemia* aff. *sinica* Cai, 1989, with a high performance of sequencing. The new primer set worked well also for other *Artemia* bisexual species, as well as for parthenogenetic populations. It is recommended that joint regions between the previous/next gene(s) and the target marker, could be aimed at when designing specific primers for other markers and taxa.

## INTRODUCTION AND METHODS

The genus *Artemia* Leach, 1819 (Branchiopoda, Anostraca), that comprises a unique taxon of zooplankton organisms, is the most conspicuous inhabitant of hypersaline environments, with a world-wide geographical distribution except on Antarctica (Zheng & Sun, 2013). The genus *Artemia* contains both bisexual species and a large number of parthenogenetic populations with different ploidy levels (Asem et al., 2010-2016a, b; Eimanifar et al., 2014).

In an ongoing comprehensive project on the phylogeography and population genetics of *Artemia* in the provinces Hebei and Qinghai (China), we have surveyed the biological communities of several inland salt lakes, especially those hosting *Artemia* aff. *sinica*. The mitochondrial *COI* (mt*COI*) marker has been utilized to identify the taxonomic status and population genetic structure, using a universal pair of primers for metazoan invertebrates (Folmer et al., 1994) and two specific primers designed especially for *Artemia* (Munoz et al., 2008; Xu, 2018). Although, these three primer pairs have been successfully amplified, the barcoding region of mt*COI* in our previous studies on different species/populations of *Artemia*, in the current project, the PCR failed for 50% to 60% of the samples examined. On the other hand, almost all successful PCR products represented poly-peak or incomplete sequences that could not be referred to a reliable sequence of mt*COI* for any phylogenetic study (figs. 1 & 2). Previously, we had made this observation in mt*COI* sequences of the pentaploid parthenogenetic population from the Yinggehai Saltern (China). The same challenges have also been noted for the Mongolian bisexual *Artemia* aff. *sinica* (S. C. Sun, pers. comm.) as well as for the majority of parthenogenetic populations from inland salt lakes in Rumania (A. Berindean, pers. comm.), in which the PCR products were not completely sequenced.

**Fig. 1.**
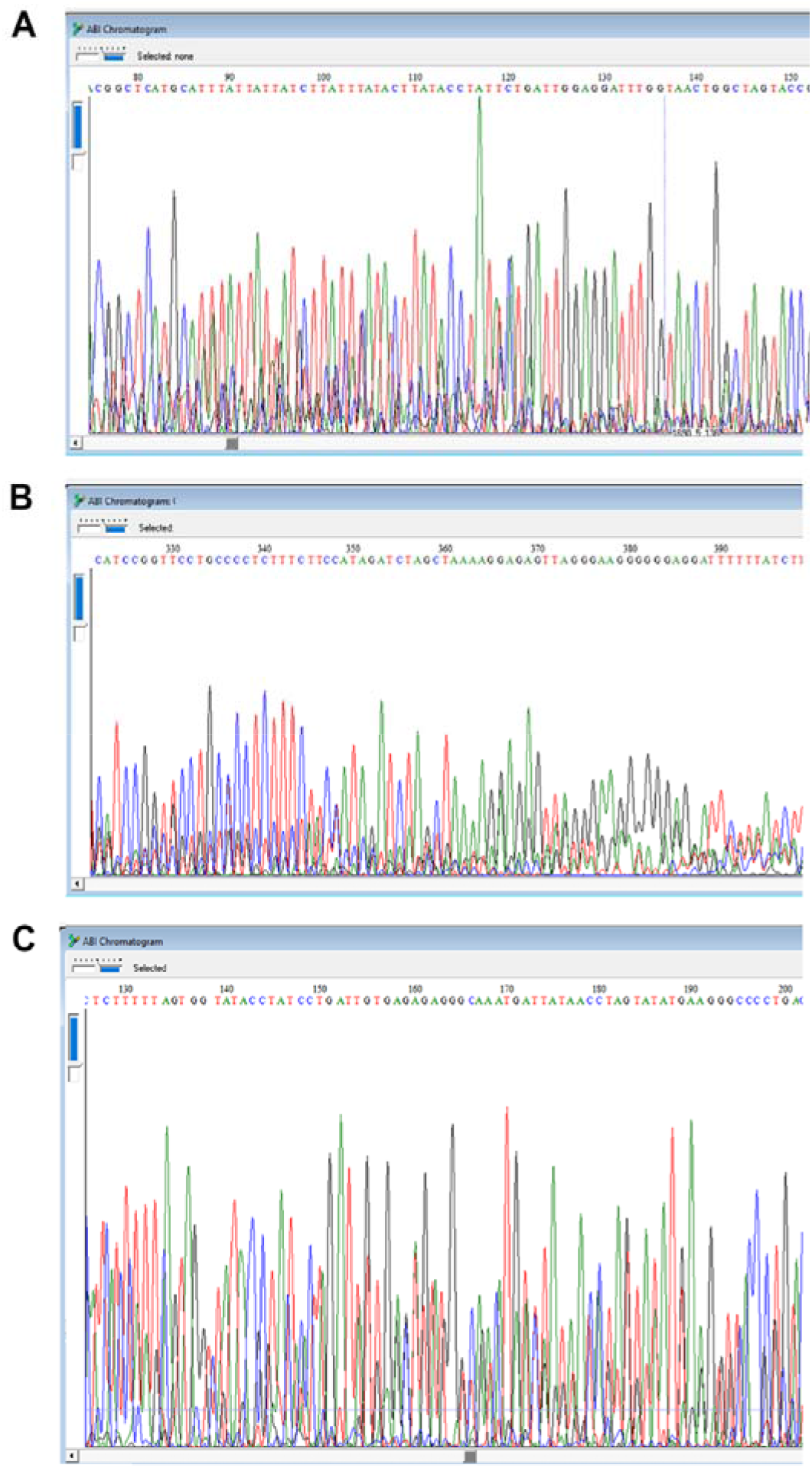
Examples of sequencing results of *Artemia* aff. *sinica* using different primers: A, Folmer et al., 1994; B, Munoz et al., 2008; C, Xu, 2018.

**Fig. 2.**
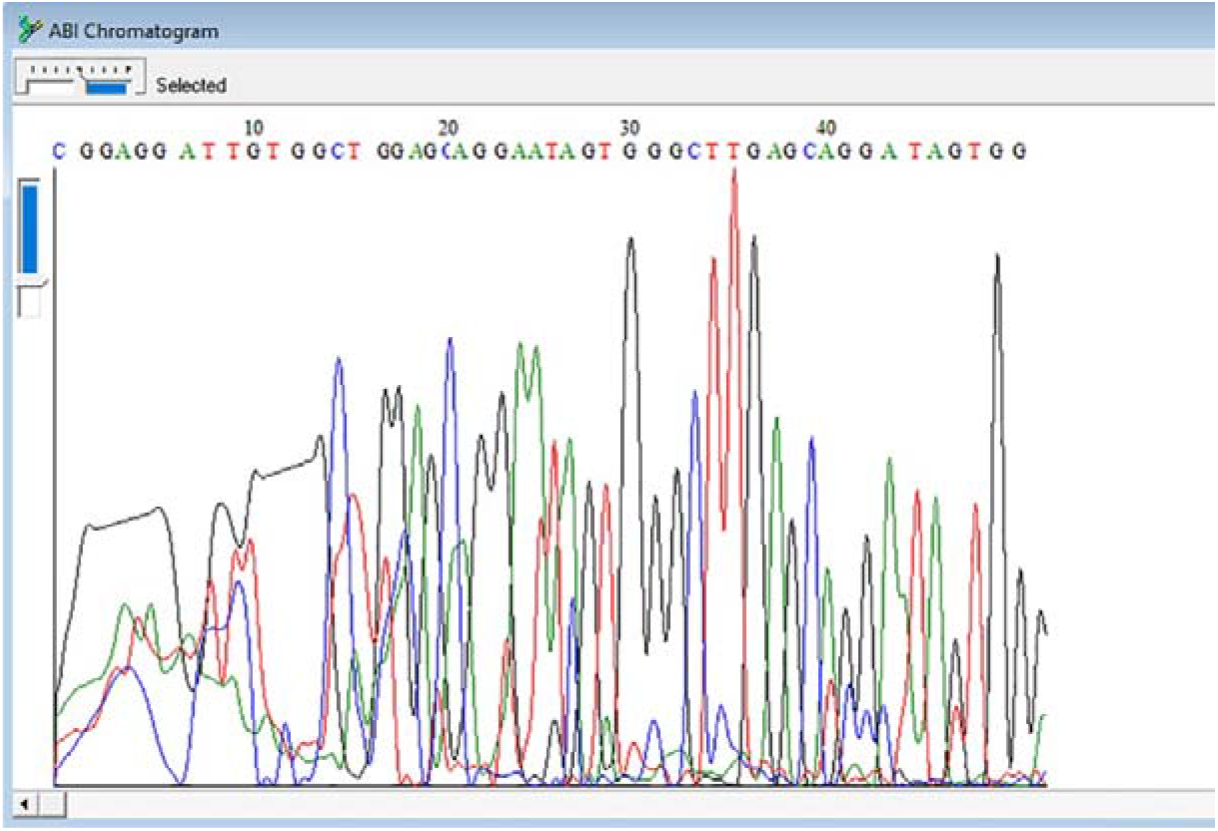
An example of incomplete sequence of *Artemia* aff. *sinica* for amplified *COI* marker with successfully <700 pb length on electrophoreses gel using previous primers (Folmer et al., 1994; B, Munoz et al., 2008; C, Xu, 2018).

Recently, a bisexual population from Mongolia has been named *Artemia frameshifta* Naganawa & Mura, 2017, using a single sequence of the mt*COI* gene (LC195588) without any confirmation from either population genetic, or biological analyses. A main problem determined in the submitted *COI* sequence, was the identification of several stop codons in the corresponding protein translation. Overall, due to those reasons, the taxonomic status of the form named *A. frameshifta* could not be confirmed (Asem et al., 2020; Eimanifar et al., 2020; D. C. Rogers, pers. comm.). Additionally, Wang et al. (2008) have reported poly peaks sequences for mt*COI* marker of *Artemia* population from Co Qen in Tibet (China). This matter might be attributed to nuclear copy/ies of mtDNA gene(s) (Srinivasainagendra et al., 2017; Eimanifar et al., 2020) and/or the existence of pseudogene(s).

In order to solve the problem, we aimed at amplifying the assured mt*COI* barcoding region of *Artemia* aff. *sinica.*We hypothesized that, if a forward primer could be designed to start amplification from the joint regions of the last part of the previous gene (tRNA*_Tyr_*) and the beginning of the target mt*COI*, the obtained sequence can most probably be referred to an errorless sequence of the mitogenomic *COI* (fig. 3).

**Fig. 3.**
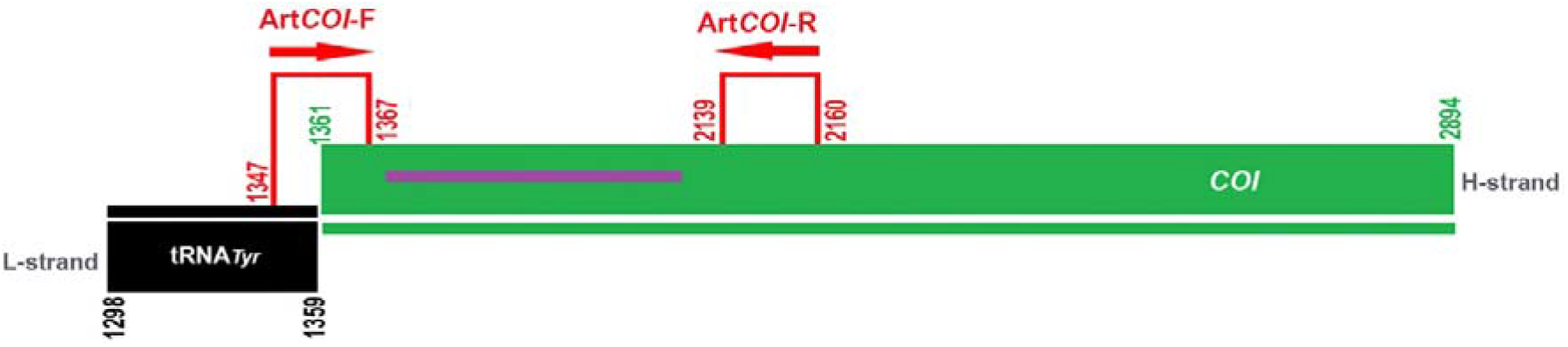
Schematic arrangement of the tRNA*_Tyr_*/*COI* positions and the relative regions of the newly designed F and R primers. The numbers show the positions on the *Artemia sinica* Cai, 1989 mitogenome. The purple line represents the *COI* barcoding region in metazoan invertebrates (to represent clear positions in this schematic figure, actual scales were not considered).

## RESULTS AND DISCUSSION

The complete mitochondrial genome of *Artemia sinica* (MK069595) was chosen as a reference sequence (Asem et al., 2019). The online primer designing tool of NCBI was employed to design the desired primers (NCBI, 2021-2022). According to the gene order of the *Artemia* mitogenome, tRNA*_Tyr_* is placed at the rear of *COI*. Because tRNA*_Tyr_* is located on the L-strand (Asem et al., 2019, 2021a), its reverse sequence on the H-strand was followed to design the F-primer. In practice this meant, that the last 13 bp of the reverse sequence of tRNA_*Tyr*_ on the H-strand (1347-1359 sites) and 7 bp of the beginning of *COI* (1361-1367 sites) were chosen to design the F-primer. Then, based on the barcoding region of the *COI* marker (Folmer et al., 1994), a fragment of 21 bp length was chosen on the mitogenome of *A. sinica* (MK069595) from 2139 to 2160 sites, to design the R-primer (fig. 3). This novel primer set:

Art*COI*-F (5 □-CGGCCACTTTACTATGCAACG-3 □) and
Art*COI*-R (5 □-CCGAATGCTTCCTTTTTCCCTC-3 □)

was designed for the barcoding region of *COI*. A fragment of 750 bp was amplified using the following PCR programme: 4 min. at 95°C, followed by 35 cycles of 60 s at 95°C, 60 s at 57-60°C and 90 s at 72°C, with a final extension step of 7 min. at 72°C. The performance of the newly designed primer set was examined for two populations of *Artemia* aff. *sinica* from China, as well as for all bisexual species and parthenogenetic populations with different ploidy levels. For parthenogenetic populations, offspring of the second generation were used to extract DNA and check ploidy levels.

The designed primers were tested and it appeared that they could successfully amplify the *COI* target region in *Artemia* aff. *sinica*, with a high quality performance in sequencing specially R reaction (fig. 4). This primer set has also exhibited its efficiency in amplification of the *COI* marker in other bisexual species and in parthenogenetic populations of *Artemia.*The list of studied species and populations has been summarized in table I. We thus designed the forward and reverse primers, Art *COI*-F and Art*COI*-R, that completely cover the *COI* barcoding region for metazoan invertebrates (Folmer et al., 1994) (see fig. 3).

**Fig. 4.**
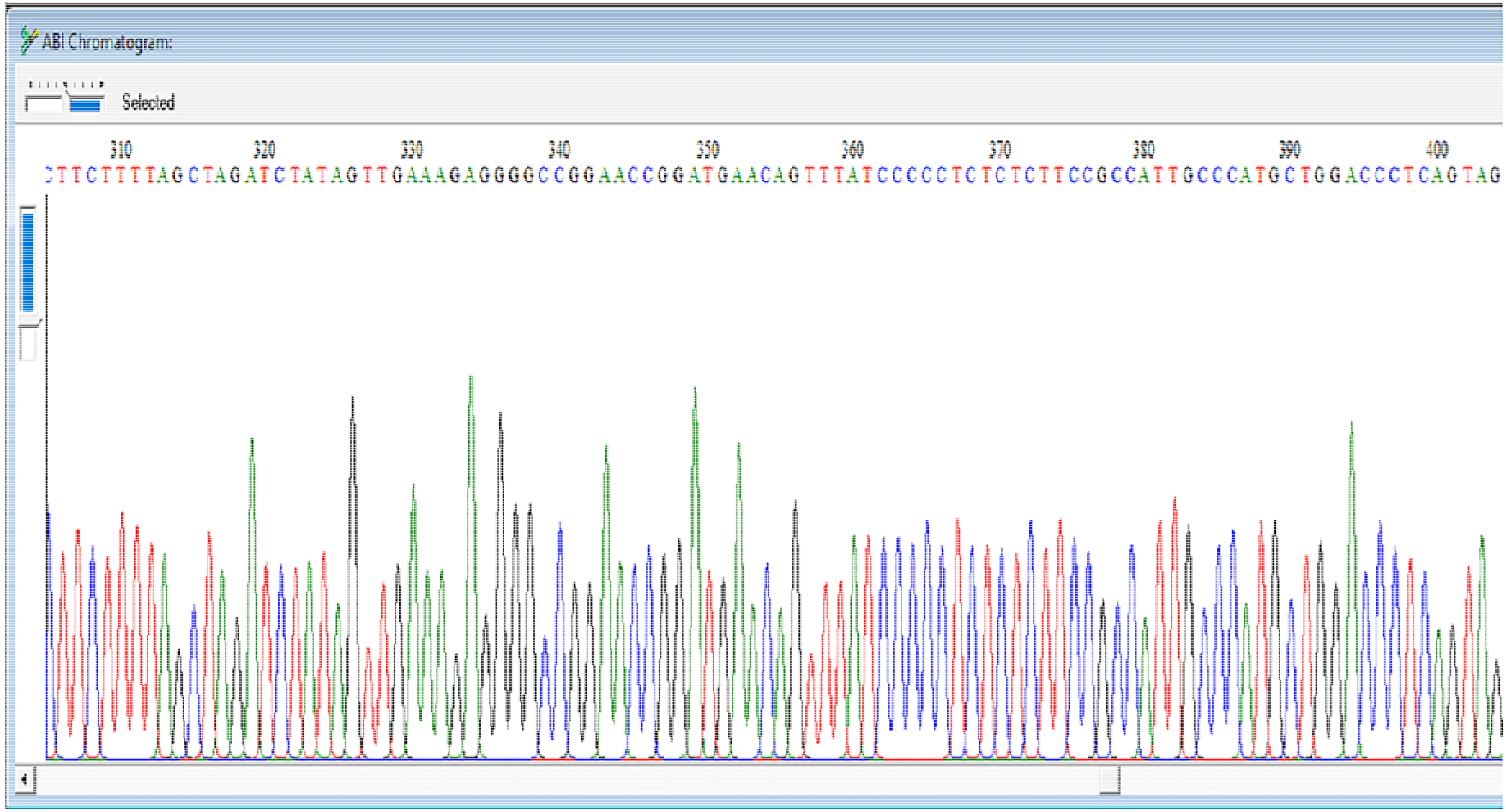
An example of sequencing results of *Artemia* aff *sinica* using the new primer set (Art*COI*-F/Art*COI*-R) described herein.

**Table I.** *Artemia* species/populations tested using Art*COI Artemia*-specific primers

| Species/population | Locality | GenBank<br>accession numbers |
| --- | --- | --- |
| *Artemia sinica* Cai, 1989 | Yuncheng Lake, China | OM737895-99 |
| *Artemia* aff. *sinica* | Hebei, China | OM737900-04 |
| *Artemia* aff. *sinica* | Qinghai, China | OM737905-09 |
| *Artemia tibetiana* Abatzopoulos, Zhang & Sorgeloos, 1998 | Lagkor Co, China | OM737910-14 |
| *Artemia urmiana* Gunther, 1889 | Urmia lake, Iran | OM737915-19 |
| *Artemia salina* (Linnaeus, 1758) | Wadi Natron, Egypt | OM737920-24 |
| *Artemia persimilis* Piccinelli & Prosdocimi, 1968 | ARC1321, Argentina | OM737925-29 |
| *Artemia franciscana* Kellogg, 1906 | Great Salt Lake, U.S.A. | OM737930-34 |
| Parthenogenetic pop. (2n) | Ga Hai Lake, China | OM737935-39 |
| Parthenogenetic pop. (3n) | Ankiembe, Madagascar | OM737940-44 |
| Parthenogenetic pop. (4n) | Hoh Lake, China | OM737945-49 |
| Parthenogenetic pop. (5n) | Yinggehai Saltern, China | OM737950-54 |

Due to the lack of unequivocal morphological characters, there is no classic identification key for *Artemia* species (Asem et al, 2010-2020). Therefore, molecular phylogenetic methods have been employed to delimit the taxonomic status of *Artemia* populations (Munoz et al., 2008; Maccari et al., 2013; Eimanifar et al., 2014, 2020; Asem et al., 2016b-2019-2021b; Saji et al., 2019; Shen et al., 2021). Hence, it is necessary to provide a dataset of a large number of sequences from each population, to reliably analyse population genetic and phylogenetic relationships among them (Maccari et al., 2013). While all primers hitherto suggested for the *COI* marker have-failed, the new primer set we developed has presented consistent and most promising results for *Artemia* aff. *sinica*.

In conclusion, we may state that, as all bisexual species and parthenogenetic populations tested in this study could be successfully sequenced in regard of their *COI* mitochondrion marker, the primer set Art*COI*-F/Art*COI*-R definitely constitutes a useful tool in phylogenetic studies of *Artemia.* It is therefore suggested in general that joint regions between the previous/next gene(s) and the target marker, be explicitly considered when designing specific primers for other markers and taxa.

## ACKNOWLEDGEMENTS

The authors thank Prof. Shi-Chun Sun (Ocean University of China, China) and Prof. Gilbert Van Stappen (*Artemia* Research Center, Ghent University, Belgium) for preparing *Artemia* cyst samples. This study has been supported by 2021 Hebei Province introduced foreign intelligence project.

## Notes

### Competing Interest Statement

The authors have declared no competing interest.

